# BCL-3 enhances β-catenin signalling in colorectal tumour cells promoting a cancer stem cell phenotype

**DOI:** 10.1101/178004

**Authors:** Danny N Legge, Alex P Shephard, Tracey J Collard, Alexander Greenhough, Adam C Chambers, Richard W Clarkson, Christos Paraskeva, Ann C Williams

## Abstract

Increased nuclear BCL-3 (a key regulator of inflammation and NF-κB signalling when associated with p50 or p52 homodimers) has been reported in a subset of colorectal cancers, but its role in colorectal tumorigenesis remains poorly understood. Interestingly, recent studies have highlighted the importance of the interplay between NF-κB signalling and the Wnt/β-catenin pathway in colorectal epithelial cells, reporting that non-stem cells engineered to undergo high levels of Wnt and NF-κB signalling can de-differentiate, initiating tumours in mice. Here we show that BCL-3 is an important co-activator of β-catenin/TCF-mediated transcriptional activity in colorectal cancer cells, increasing expression of Wnt-regulated intestinal stem cell genes. We demonstrate RNAi-mediated BCL-3 suppression reduced β-catenin/TCF-dependent transcription and the expression of intestinal stem cell genes and Wnt targets *LGR5* and *ASCL2*. Further we show that BCL-3 promotes the stem cell phenotype in colorectal cancer cells by increasing colorectal spheroid and tumoursphere formation in 3D culture conditions. Our data suggest that targeting BCL-3 may represent an exciting new approach for CRC treatment, particularly as it acts downstream of frequently mutated APC and β-catenin.

## Introduction

Around 49000 deaths per year are attributed to colorectal cancer (CRC) in the United States alone and in 2016 it’s estimated that the number of CRC-related deaths will be second only to lung cancer [1]. The B-Cell Chronic Lymphocytic Leukaemia/Lymphoma 3 (BCL-3) protein is highly expressed in a subset of CRCs where we have recently shown it inhibits apoptosis and promotes tumour growth [2]. BCL-3 is an atypical member of the inhibitor of kappa B (IκB) family of proteins and has been demonstrated to modulate transcription of NF-κB target genes via binding to homo-dimeric subunits of p50 or p52 through its ankyrin-repeat domains [3, 4]. The p50/p52 subunits possess DNA-binding motifs enabling them to occupy κB sites at promoters of NF-κB-responsive genes [5], permitting BCL-3 to activate (through its own transactivation domain or via recruiting alternative co-activators) or repress gene transcription [6].

Although first characterised in hematopoietic cancers [7], there is an emerging role for BCL-3 in solid tumours. It has been implicated in cancers arising in multiple tissue types including breast[8], prostate[9], cervical [10] and colorectal [2]. BCL-3 bears numerous tumour-promoting capabilities such as increasing proliferation [11], inflammation [12], inhibiting apoptosis [13] and promoting metastasis [14]. Importantly, studies by Puvvada *et al.* [15] and more recently by Saamarthy *et al.* [16] report that around 30% of colorectal tumours present with nuclear BCL-3.

The Wnt pathway plays a critical role in gut development, maintenance and homeostasis [17]. During active signalling, the Wnt effector protein β-catenin translocates from the cytoplasm to the nucleus and binds TCF/LEF transcription factors situated at Wnt-responsive genes. β-catenin recruits other co-activators such as CBP [18] and BCL-9 [19] to initiate transcription of genes involved in proliferation or stemness that are otherwise silent in the absence of Wnt ligands [20]. Co-activators that bind to the C-terminus of β-catenin are diverse in their methods of transcriptional activation and include chromatin-remodelling enzymes, histone acetyl-transferases and histone methyl-transferases [20]. CRCs frequently occur through inactivating mutations of the tumour suppressor APC (part of the ‘destruction complex’ that degrades β-catenin) or less-frequently via stabilizing mutations in β-catenin itself [21], consequently resulting in deregulated β-catenin signalling.

Considering the importance of Wnt/β-catenin and NF-κB crosstalk in cellular de-differentiation and tumour initiation in the intestine [22], surprisingly there have been no studies exploring the role of NF-κB co-regulator BCL-3 in Wnt signalling in CRC. Here we report that BCL-3 is an important co-activator of β-catenin/TCF-mediated transcriptional activity in CRC cells. We show that BCL-3 regulates β-catenin-mediated transcription and expression of colorectal stem cell and cancer stem cell marker genes *LGR5* and *ASCL2*, promoting a stem cell phenotype in cancer cells. The data we present in this study suggest BCL-3 is important in promoting a cancer stem cell phenotype and therefore could be an attractive target for future therapies.

## Results and Discussion

### **BCL-3 interacts with** β**-catenin and modulates** β**-catenin/TCF reporter activity**

To examine the role of BCL-3 in Wnt/β-catenin signalling initially we screened a panel of human adenoma and carcinoma-derived cell lines for expression of BCL-3, β-catenin, TCF4 and LEF1 (Supplementary Figure 1). To investigate any potential interaction between BCL-3 and β-catenin in CRC cells we selected the *APC* mutant SW1463 cell line for its high endogenous expression of both BCL-3 and β-catenin. We performed BCL-3 Co-IPs on nuclear-enriched lysates and were able to identify an interaction between endogenous BCL-3 and β-catenin previously unreported in CRC cells (Figure 1A). It has been demonstrated that TNF-α can induce BCL-3 binding to p52 homodimers [23], therefore as a positive control we treated SW1463 cells with TNF-α for 6 hours to activate NF-κB signalling and carried out BCL-3 Co-IPs in the resulting lysates. In stimulated cells we detected an interaction between BCL-3 and p52 and again demonstrated the association of BCL-3 and β-catenin (Figure 1B). These data suggest endogenous BCL-3 interacts with β-catenin in CRC cells.

**Figure 1.**
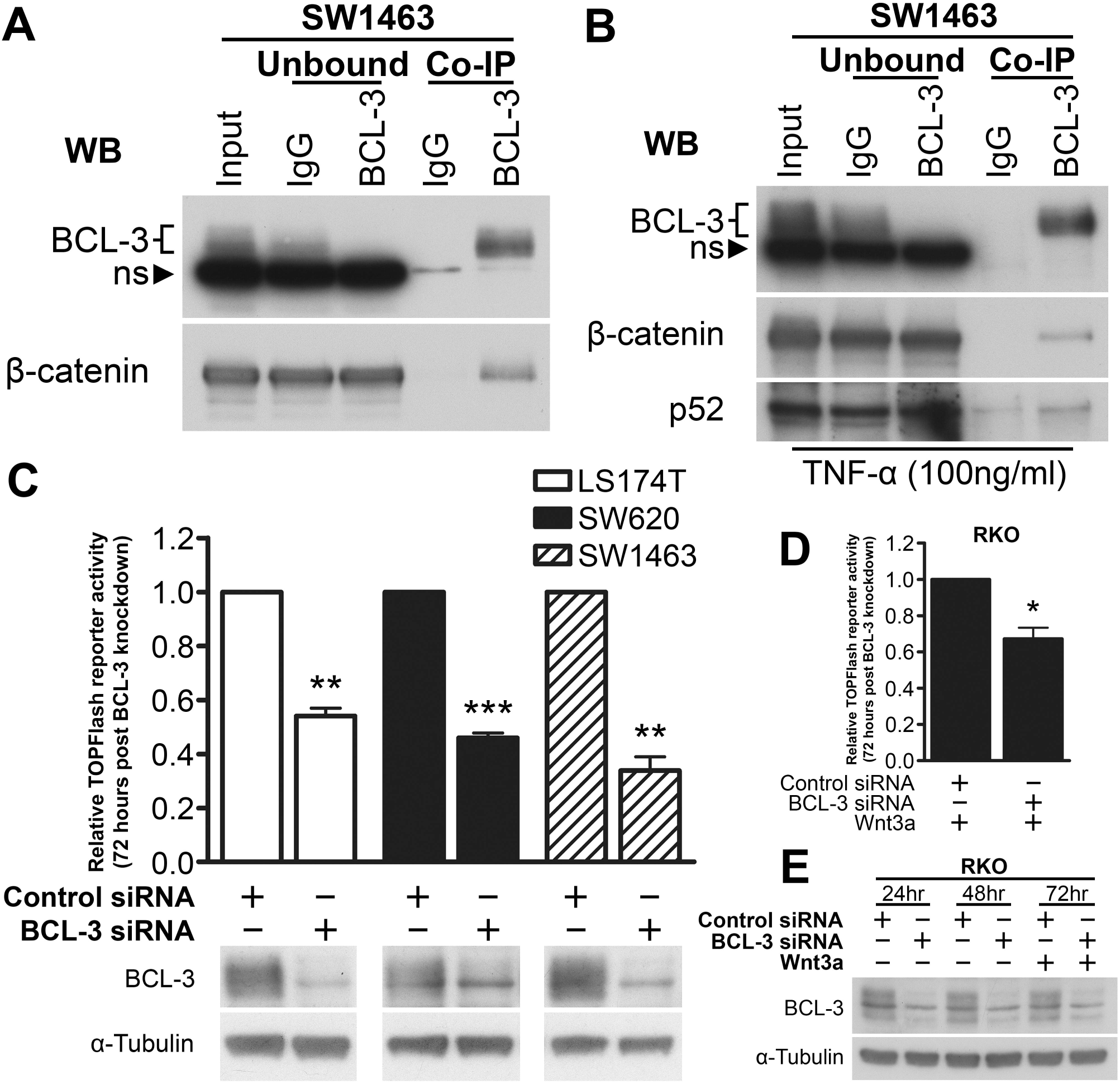
BCL-3 interacts with β-catenin and modulates β-catenin/TCF reporter activity in colorectal cancer cells. (**A**) and (**B**) BCL-3 Complex-immunoprecipitation (Co-IP) performed in SW1463 nuclear-enriched lysates. Unbound (Immuno-depleted lysates) shows depletion of proteins after IP. Immunoprecipitates were analysed by western blot for BCL-3 and β-catenin. IgG serves as negative control. (**B**) Cells were treated with 100ng/ml TNF-α for 6 hours prior to lysis. Immunoprecipitates were additionally analysed for p52. Note the presence of non-specific band in BCL-3 western analysis with use of Abcam BCL-3 antibody. **Unbound** = immuno-depleted lysate. **ns** = non-specific band. **WB** = western blot antibody. (**C**) β-catenin/TCF (TOPFlash) luciferase reporter assay. TOPFlash reporter activity measured 72 hours post-siRNA transfection in LS174T, SW620 and SW1463 carcinoma cells. Results are expressed as TOPFlash/Renilla. Western analysis shows expression of BCL-3. α-Tubulin serves as a loading control. (**D**) TOPFlash reporter activity measured 72 hours post-siRNA transfection in RKO cells. 100ng/ml Wnt3a was added to cells at 48 hours post-siRNA transfection. Results are expressed as TOPFlash/FOPFlash. One sample t-test. *N*=3, ± SEM \**P*<0.05, \*\**P*<0.01, \*\*\**P*<0.001. (**E**) Western blot analysis of BCL-3 expression over duration of TOPFlash assay performed in (**D**). α-Tubulin serves as a loading control.

We next investigated the effects of BCL-3 expression on β-catenin/TCF-mediated transcription. To do this we used siRNA to suppress *BCL3* expression in colorectal cell lines before transfecting cells with TOPFlash reporter plasmid to measure β-catenin/TCF-mediated transcriptional output. Interestingly, we discovered a significant decrease in TOPFlash activity in LS174T (colon-derived, mutant β-catenin), SW620 (lymph node-derived, mutant APC) and SW1463 (rectal-derived, mutant APC) cell lines (Figure 1C). These data indicate that BCL-3 could promote β-catenin/TCF-mediated transcription in CRCs with common Wnt driver mutations. In addition, we examined the role of BCL-3 in RKO CRC cells, which are reported to harbour no activating Wnt pathway mutations and show no detectable TOPFlash activity under unstimulated conditions [24]. In agreement with preceding experiments, there was a significant decrease in Wnt3a-induced TOPFlash activity in RKO cells depleted for BCL-3 (Figure 1D). These findings show that in a non-deregulated Wnt setting BCL-3 can modulate β-catenin/TCF-dependent transcription, suggesting Wnt3a-mediated transcriptional responses are enhanced by BCL-3.

### BCL-3 regulates stemness-associated Wnt targets

After establishing that BCL-3 regulates β-catenin/TCF-dependent transcription, we investigated the role of BCL-3 in Wnt target gene regulation. Wnt targets are heavily involved in the maintenance of stem cells in the colon [17], with *LGR5* and *ASCL2* providing key examples of genes that are expressed in intestinal stem cells, but not expressed in other cell types of the gut [25-28]. With this in mind, we used qRT-PCR to analyse mRNA levels of some classical Wnt target genes—along with stemness-associated Wnt targets—upon BCL-3 suppression in LS174T, SW620 and SW1463 cells. We found classical β-catenin/TCF targets such as c-MYC and Cyclin D1 were not significantly regulated by BCL-3 knockdown, whereas stem cell-specific Wnt targets *LGR5* and *ASCL2* were significantly downregulated in all three cell lines (Figure 2A). Western analysis was used to measure LGR5 and ASCL2 protein expression at 48 and 72 hours post-BCL-3 siRNA transfection; we found clear downregulation of LGR5 and ASCL2 protein in all cell lines and at both time-points (Figure 2B). Interestingly, we saw no decrease in total or ‘actively signalling’ (de-phosphorylated) β-catenin levels. Furthermore, using LS174T and SW620 cell lines stably-expressing a BCL-3 expression construct (termed LS174T^BCL-3^ and SW620^BCL-3^) or empty vector (LS174T^pcDNA^ and SW620^pcDNA^) we were able to show increased expression of BCL-3 enhanced LGR5 expression (Supplementary Figure 2). Together, these data suggest that BCL-3 may be promoting β-catenin/TCF-dependent transcription at specific intestinal stemness-associated gene loci, which doesn’t appear to be via increasing the pool of actively signalling β-catenin.

**Figure 2.**
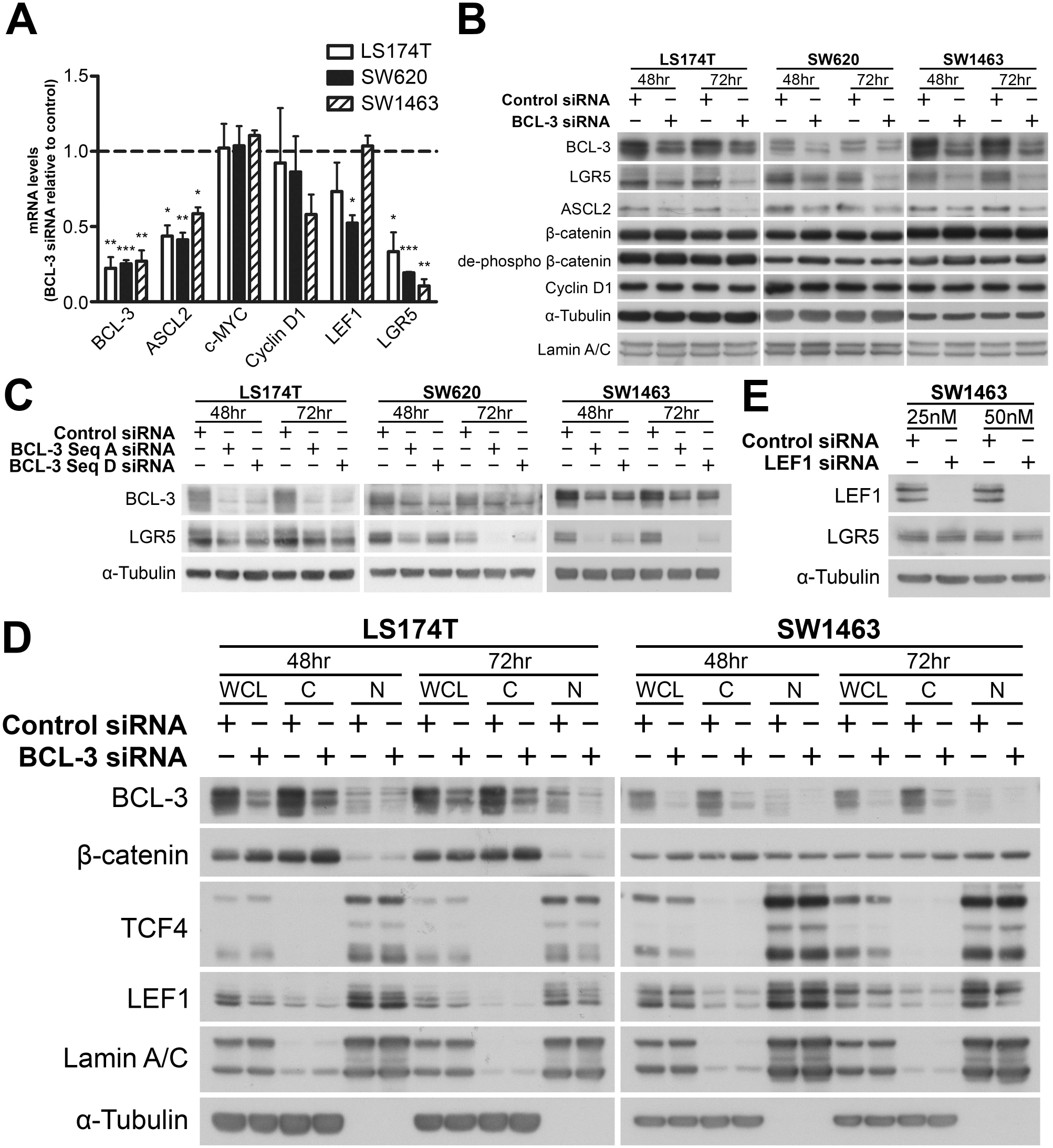
BCL-3 regulates expression of intestinal stem cell markers and does not promote nuclear translocation of β-catenin. (**A**) Quantitative reverse transcriptase-PCR (qRT-PCR) mRNA analysis of Wnt target gene expression in LS174T, SW620 and SW1463 cells following BCL-3 suppression. Data shows mRNA expression of genes normalised to housekeeping genes at 72 hours post-siRNA transfection, BCL-3 knockdown cells relative to controls. One sample t-test. *N*=3, ± SEM \**P*<0.05, \*\**P*<0.01, \*\*\**P*<0.001. (**B**) Western analysis of LGR5 and ASCL2 expression in LS174T, SW620 and SW1463 cells following BCL-3 suppression. Expression of total β-catenin, de-phosphorylated (actively signalling) β-catenin and Cyclin D1 also shown. α-Tubulin serves as a loading control. ASCL2 expression is shown in nuclear-enriched lysates, with Lamin A/C serving as a loading control. (**C**) LGR5 expression in LS174T, SW620 and SW1463 cells following BCL-3 suppression with two independent siRNA sequences. LGR5 and BCL-3 expression analysed by western blot. α-Tubulin serves as a loading control. (**D**) Nuclear/cytoplasmic enrichments of LS174T and SW1463 cells following BCL-3 suppression showing localisation and expression of Wnt transcriptional regulators β-catenin, TCF4 and LEF1. Western analysis confirms BCL-3 suppression. α-Tubulin and Lamin A/C serve as loading controls and confirm cytoplasmic/nuclear enrichment. **WCL** = whole cell lysate. **C** = cytoplasm-enriched lysate. **N** = nuclear-enriched lysate. (**E**) LGR5 expression following LEF1 suppression with 25nM and 50nM siRNA in SW1463 cells. Data shown 48 hours post-siRNA transfection. α-Tubulin serves as a loading control.

*LGR5* encodes a G-protein coupled receptor expressed in stem cells of the intestine and is also thought to identify cancer stem cells in CRC [27, 29-31]. To further confirm regulation of LGR5 by BCL-3—and to rule out any potential off-target effects of using a single siRNA sequence—we used a second BCL-3-targeting siRNA. LGR5 suppression by BCL-3 depletion was demonstrated using two independent siRNA sequences in SW620, LS174T and SW1463 cell lines (Figure 2C), indicating that BCL-3 regulates LGR5 expression in CRC cells of different origins and mutational backgrounds.

By preferentially diverting β-catenin/TCF-mediated transcription towards stemness genes *LGR5* and *ASCL2* over some of the more classical Wnt targets such as Cyclin D1 and c-MYC—as recently reported for R-Spondin 3 translocations in CRC [32]—it appears BCL-3 may be driving CRC cells towards a more stem-cell phenotype. Not only are *LGR5* and *ASCL2* robust stem cell markers but numerous studies have shown them to be upregulated in CRCs, with some reporting they are markers for CRC stem cells [29, 30, 33-36].

### **BCL-3 does not mediate** β**-catenin activity through promoting nuclear translocation or altering levels of LEF1**

Having shown nuclear interaction between BCL-3 and β-catenin and given that BCL-3 has been shown to promote NF-κB nuclear translocation [37], to further examine how BCL-3 regulates β-catenin/TCF activity and target gene expression we investigated whether BCL-3 enhances β-catenin nuclear localisation. We performed western analysis on LS174T and SW1463 nuclear/cytoplasmic-enriched lysates following BCL-3 suppression. BCL-3 knockdown did not affect β-catenin nuclear translocation in either cell line (Figure 2D). We also analysed expression levels of TCF4 and LEF1, as diminished nuclear levels of these transcription factors may have a profound effect on transcription of Wnt target genes. We found no consistent regulation of TCF4, but a small decrease in LEF1 expression (Figure 2D). *LEF1* is a Wnt target gene and encodes a transcription factor that mediates β-catenin signalling [38, 39]. To investigate the role of LEF1 in BCL-3-mediated LGR5 regulation we used RNAi to suppress LEF1 in SW1463 cells and analysed LGR5 expression using western blotting. We found LEF1 knockdown had no effect on LGR5 levels (Figure 2E). These findings signify that modulation of β-catenin/TCF-dependent transcription and regulation of stemness-associated Wnt targets by BCL-3 is not through promotion of β-catenin, TCF4 or LEF1 nuclear translocation. When taken alongside our data from Co-IP and TOPFlash experiments (Figure 1) these results support the idea that BCL-3 may be acting as a transcriptional co-activator of the β-catenin/TCF complex.

### BCL-3 regulates colorectal spheroid and tumoursphere formation in 3D culture

After demonstrating that BCL-3 promotes LGR5 and ASCL2 expression, we investigated the functional consequence of stemness marker regulation using an adapted version of the organoid model system pioneered by Sato *et al.* [25] to grow CRC cell lines as 3D spheroids. 3D culture provides the additional benefit of creating more relevant physiological conditions to study tumour growth and initiation when compared with 2D cell culture. It has been shown that single LGR5-positive cells can form organoids in Matrigel [25, 40]. As BCL-3 promotes LGR5 expression, we hypothesised that suppressing BCL-3 may inhibit the ability of single cells to progress and form spheroids. To test this, we transfected SW1463 cells with control or BCL-3 siRNA for 24 hours. SW1463 cells were used as they form lumenal spheroids in 3D culture (Figure 3B), suggesting the presence of differentiated cell types in addition to cancer stem cells [41, 42]. Equal numbers of viable control and BCL-3 siRNA treated cells were re-suspended in Matrigel and seeded into 48 well plates. We then measured the number of spheroids formed after 10 days of culture. At the time of seeding, duplicate flasks were lysed and checked for efficacy of BCL-3 knockdown. We found suppression of BCL-3 and downregulation of LGR5 was achieved after 24 hours (Figure 3A). Furthermore, we noted a significant reduction in the number of spheroids formed upon BCL-3 suppression relative to controls after 10 days of growth (Figures 3C and 3D). We repeated this experiment using SW620^pcDNA^ and SW620^BCL-3^ cells to investigate the outcome of BCL-3 overexpression on spheroid formation. We found BCL-3 overexpression significantly increased the number of spheroids formed following 10 days of culture (Figure 3E). These results suggest BCL-3 enhances the ability of cells to initiate full spheroid formation under these conditions, indicating that BCL-3 may promote stemness of CRC cells.

**Figure 3.**
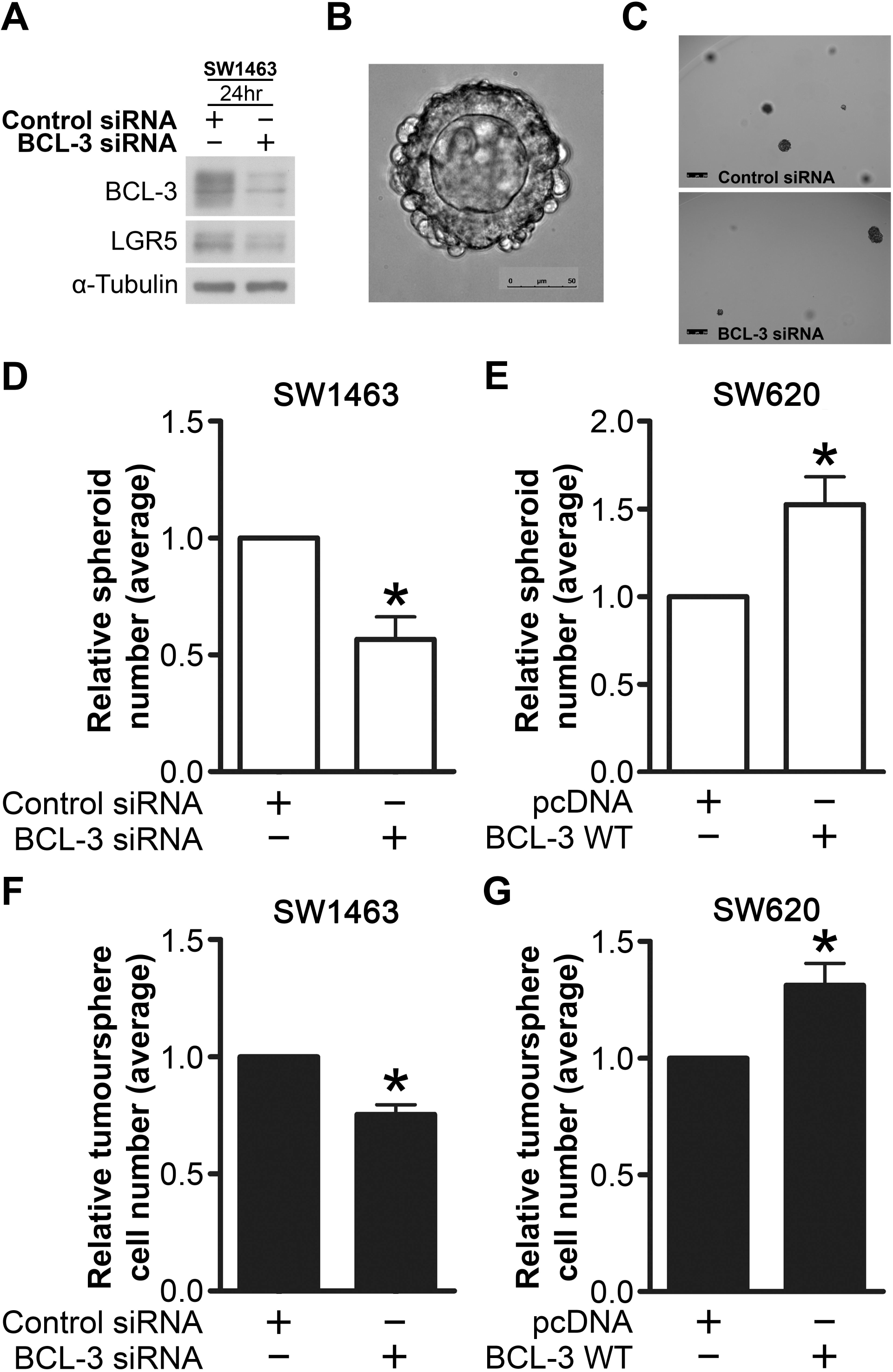
BCL-3 regulates colorectal spheroid and tumoursphere formation. (**A**) Western analysis confirming BCL-3 suppression and LGR5 downregulation in SW1463 cells seeded into Matrigel. α-Tubulin serves as a loading control. (**B**) Widefield microscopy image of a spheroid grown from a single SW1463 cell following 10 days of culture. 20x objective. Scale bar = 50μm. (**C**) Widefield microscopy images of wells containing BCL-3 knockdown and control SW1463 spheroids. 5x objective. Scale bars = 250μm. (**D**) and (**E**) Spheroid forming assay. (**D**) SW1463 cells transfected with BCL-3 siRNA or negative control siRNA were re-suspended in Matrigel and seeded into 48 well plates, 24 hours post-transfection. Spheroids were cultured for 10 days as previously described [52]. Wells were imaged and analysed using Matlab R2015a software, with subjective gating applied to exclude any cells/debris less than 3000μm^2^ in cross-sectional area. (**E**) SW620 cells stably-transfected with pcDNA-BCL-3 WT vector (SW620^BCL-3^) or empty pcDNA vector (SW620^pcDNA^) as a negative control were seeded into Matrigel, cultured and analysed as in (**D**). One sample t-test. *N*=3, ± SEM \**P*<0.05. (**F**) and (**G**) Tumoursphere forming assay. SW1463 cells transfected with BCL-3 siRNA or negative control siRNA were re-suspended in tumoursphere medium and cultured as described previously [43]. Tumourspheres were dissociated and cells counted after 6 days of culture. (**G**) SW620 cells stably-transfected with pcDNA-BCL-3 WT vector or empty pcDNA vector as a negative control were seeded, cultured and analysed as in (**F**). One sample t-test. *N*=4, ± SEM \**P*<0.05.

To further investigate the effect of BCL-3 on stemness of CRC cells we used a tumoursphere 3D culture system. Tumoursphere assays have been used previously to identify undifferentiated cancer stem cells in CRC [43]. We transfected SW1463 cells with control or BCL-3 siRNA for 24 hours before seeding into serum-free conditions in non-adherent 6 well plates and cultured for 6 days. The number of viable cells were counted at the end of this period to determine any effect BCL-3 might have on tumoursphere formation. Intriguingly, we found BCL-3 suppression significantly reduced tumoursphere formation in serum-free conditions (Figure 3F). Furthermore, upon BCL-3 overexpression in the same system using SW620^pcDNA^ and SW620^BCL-3^ lines we reported a significant increase in the number of cells comprising the tumourspheres formed by BCL-3 overexpressing cells versus controls (Figure 3G). This suggests that by enhancing their ability to form tumourspheres in 3D culture conditions BCL-3 could be promoting stemness of CRC cells.

The cancer stem cell hypothesis states there are cells within tumours capable of self-renewal, in addition to producing other heterogeneous, differentiated cell types that constitute the tumour mass [44]. Cancer stem cells fuel tumour growth and are thought to be responsible for tumour reconstitution in cases of relapse if they are not entirely eliminated by initial treatments [29, 45]. Therefore, cancer stem cells must be targeted as a means of developing more effective therapies. In this study, enhancing levels of BCL-3 in CRC cells promoted spheroid and tumoursphere formation, indicating an increase in CRC stem-like activity [43]. Similarly, BCL-3 suppression impaired the ability of single cells to progress to fully-formed spheroids and inhibited tumoursphere formation. Through using colorectal spheroid and tumoursphere forming assays we present functional evidence that BCL-3 is promoting cancer stem cell activity. Intriguingly, our findings complement those of a recent study by Chen *et al*. proposing that BCL-3 plays a key role in maintaining naïve pluripotency in mouse embryonic stem cells [46]. The findings of the Chen *et al.* study, albeit in a non-cancer background, concur with our conclusion that BCL-3 is important in promoting stemness. As yet the mechanism behind the selectivity of *LGR5* and *ASCL2* regulation over other canonical Wnt targets remains undefined and warrants further investigation. It may be governed by other co-regulators present in the transcriptional complex, as has been previously demonstrated by Kim *et al.* [47] who showed both activation and repression of *KAI1* occurred in the presence of BCL-3, with activation or repression dependent on the co-regulator Pontin or Reptin, respectively.

In conclusion, we have shown for the first time that BCL-3 potentiates β-catenin/TCF-mediated signalling in CRC and selectively regulates transcription of intestinal stem cell genes and Wnt targets *LGR5* and *ASCL2*, promoting a cancer stem cell phenotype. Our data suggest BCL-3 may represent an exciting new avenue for targeting cancer stem cells in CRC, particularly as it enhances β-catenin activity downstream of frequently mutated APC and β-catenin.

## Materials and Methods

### Cell culture

All cell lines were cultured in 10% DMEM -Dulbecco’s Modified Eagle Medium (DMEM) (Gibco, Life Technologies, Paisley, UK) with added 10% foetal bovine serum (FBS) (GE Healthcare, UK), 2mM glutamine (Sigma-Aldrich, Dorset, UK), 100 units/ml penicillin and 100 units/ml streptomycin (Invitrogen, Life Technologies, Paisley, UK). LS174T^pcDNA/BCL-3^ and SW620^pcDNA/BCL-3^ cell lines were cultured in 10% DMEM with added 400μg/ml G418 (Sigma-Aldrich). For stock purposes cells were maintained in T25 flasks (Corning, NY, USA) and incubated at 37°C in dry incubators maintained at 5% CO_2_. Cells were medium changed every 3-4 days.

### SDS-PAGE and western analysis

Cell lysates were prepared and subjected to western analysis as described previously [48] using the following antibodies: α-Tubulin (T9026; Sigma-Aldrich), ASCL2 (4418; EMD Millipore, Watford, UK), β-catenin (9587; Cell Signalling Technology, MA, USA), β-catenin (610153; BD Biosciences, CA, USA), de-phosphorylated (actively signalling) β-catenin (05-665; EMD Millipore), BCL-3 (ab49470; Abcam, Cambridge, UK), BCL-3 (23959; Proteintech, Manchester, UK), c-MYC (sc-40; Santa Cruz Biotechnology, TX, USA), Cyclin D1 (2978; Cell Signalling Technology), Lamin A/C (4200236; Sigma), LEF1 (2230; Cell Signalling Technology), LGR5 (ab75850; Abcam), p52 (05-361; EMD Millipore) and TCF4 (2569; Cell Signalling Technology).

### Complex Immunoprecipitation (Co-IP)

BCL-3 CoIPs were performed in SW1463 nuclear-enriched lysates as previously described [49]. Briefly, 500μg of rabbit pan-IgG (12-370; EMD Millipore) pre-cleared nuclear-enriched protein lysates were cleared with 6μg of BCL-3 antibody (23959; Proteintech) conjugated to Dynabeads Protein A beads (Invitrogen) before undergoing western blot analysis. Beads were retrieved using a Dyna-Mag 2 magnet (Invitrogen). Immunoprecipitates were analysed by western blot for BCL-3 (ab49470; Abcam) and β-catenin (610153; BD Biosciences). For TNF-α treated CoIPs, 100ng/ml TNF-α (Source BioScience, Nottingham, UK) was added to cells for 6 hours prior to lysis. Immunoprecipitates were additionally analysed for p52 (05-361; EMD Millipore)

### Gene knockdown via RNA interference (RNAi)

Cell lines were reverse transfected in Opti-MEM (Gibco) using RNAiMax (Invitrogen) with 50nM of siRNA, unless otherwise stated. Individual sequences and relevant non-targeting controls were used for silencing *BCL3* whereas a smart pool and non-targeting controls were used to silence *LEF1*. All siRNA sequences were produced by Dharmacon (GE Lifesciences).

### Quantitative reverse transcriptase-PCR (qRT-PCR)

Tri-Reagent (Sigma-Aldrich) was added to cells and RNAeasy mini kit (Qiagen, Limberg, Netherlands) was used according to manufacturer’s instructions to clean up RNA before synthesis of cDNA and qRT-PCR were performed as previously described [49] using the following primers: *BCL3* (QT00040040), *CTNNB1* (β-catenin) (QT01331274), *CMYC* (QT00035406), *CCND1* (Cyclin D1) (QT00495285), *LEF1* (QT00021133), *LGR5* (QT00027720). Gene expression was normalised to housekeeping gene *TBP* (QT00000721). *ASCL2* primer sequences (Sigma) were obtained from Jubb *et al.* [35] (forward 5’ – GGCACTGCTGCTCTGCTA – 3’, reverse 5’ – GTTCACGCTCCCTTGAAGA – 3’).

### TCF reporter assay (TOPFlash reporter assay)

BCL-3 expression was suppressed via RNAi and TOPFlash assay was performed as previously described [49] using Promega Dual-Luciferase Reporter Assay System (Promega, WI, USA) according to manufacturer’s instructions. Luminescence was measured at 560nm using a Modulus luminometer (Turner Biosciences, CA, USA). Co-transfection of FOPFlash reporter with mutated TCF consensus sites was used alongside TOPFlash to monitor non-specific output. In RKO cells 100ng/ml recombinant human WNT3a protein (R&D Systems, Abingdon, UK) was added to cells 48 hours post-siRNA transfection.

### Generation of stable BCL-3 expressing cell lines

Cells were transfected in Opti-MEM (Gibco) using Lipofectamine 2000 (Invitrogen) with pcDNA3-BCL-3 WT vector [50, 51] (kindly provided by Alain Chariot, University of Liège, Belgium; re-cloned by Tracey Collard, University of Bristol, UK) or empty pcDNA3 vector as a control. Resistant clones were selected and pooled in media supplemented with 500μg/ml neomycin.

### Spheroid forming assay

For BCL-3 knockdown experiments equal numbers of single SW1463 cells transfected with BCL-3 siRNA or negative control siRNA were re-suspended in Matrigel (Becton Dickinson, Oxford, UK) and seeded into 48 well plates (Corning), 24 hours post-transfection. Matrigel was submerged in spheroid medium consisting of Advanced DMEM:F12 (Gibco), 0.1% BSA (Sigma-Aldrich), 2mM glutamine (Sigma-Aldrich), 10mM HEPES (Sigma-Aldrich), 100 units/ml penicillin, 100 units/ml streptomycin, 1% N2 (Gibco), 2% B27 (Gibco) and 0.2% N-acetyl-cysteine (Gibco). Spheroids were cultured for 10 days as previously described [52]. Wells were imaged using a DMI6000 widefield microscope (Leica Microsystems, Wetzlar, Germany) and Leica LAS-X acquisition software (Leica Microsystems).

Images were analysed using Matlab R2015a software (Mathworks, MA, USA). Subjective gating was set to exclude any cells/debris less than 3000μm^2^ in cross-sectional area. For BCL-3 overexpression experiments equal numbers of single SW620 cells stably-transfected with pcDNA-BCL-3 WT vector or empty pcDNA vector as a negative control were seeded into Matrigel and were cultured and analysed as described above.

### Tumoursphere forming assay

For BCL-3 knockdown experiments SW1463 cells transfected with BCL-3 siRNA or negative control siRNA were counted using trypan blue (Invitrogen) and a Countess automated cell counter (Invitrogen) to exclude dead cells. 2x10^4^ viable cells were re-suspended in tumoursphere medium consisting of Dulbecco’s Modified Eagle Medium:F12 medium (DMEM:F12) (Gibco) supplemented with 20ng/ml EGF (Sigma-Aldrich), 10ng/ml FGF (R&D Systems), 2% B27 (Gibco), 400μg/ml G418, 2mM glutamine, 100 units/ml penicillin and 100 units/ml streptomycin (Invitrogen). Cells were seeded into non-adherent 6 well plates (Greiner Bio-One, Stonehouse, UK) and tumoursphere were cultured as described previously [43] for 6 days before contents of wells were centrifuged for 3 minutes at 3000rpm. Tumourspheres were re-suspended in 500μL of StemPro Accutase cell dissociation reagent (Gibco) and manually dissociated via pipetting. Cells were incubated for 30 minutes to achieve single cell suspension and viable cells were counted using a Countess cell counter. For BCL-3 overexpression experiments SW620 cells stably-transfected with pcDNA-BCL-3 WT vector or empty pcDNA vector as a negative control were seeded, cultured and analysed as above.

## Author Contributions

Conception and design [ACW, DL, AS, TJC, CP, AG, RWC], execution [ACW, DL, AS, TJC, AG, ACC], analysis and interpretation of data [ACW, DL, AS, TJC, CP, AG, ACC] drafting the article [DL, AG, ACW] revising it critically for important intellectual content [All].

## Acknowledgements

Thanks to the Wolfson Bioimaging Facility (University of Bristol) for aiding widefield microscopy experiments.

### Conflict of Interest

The authors declare no conflict of interest.

## Supplementary figure legends

**Supplementary Figure 1.**
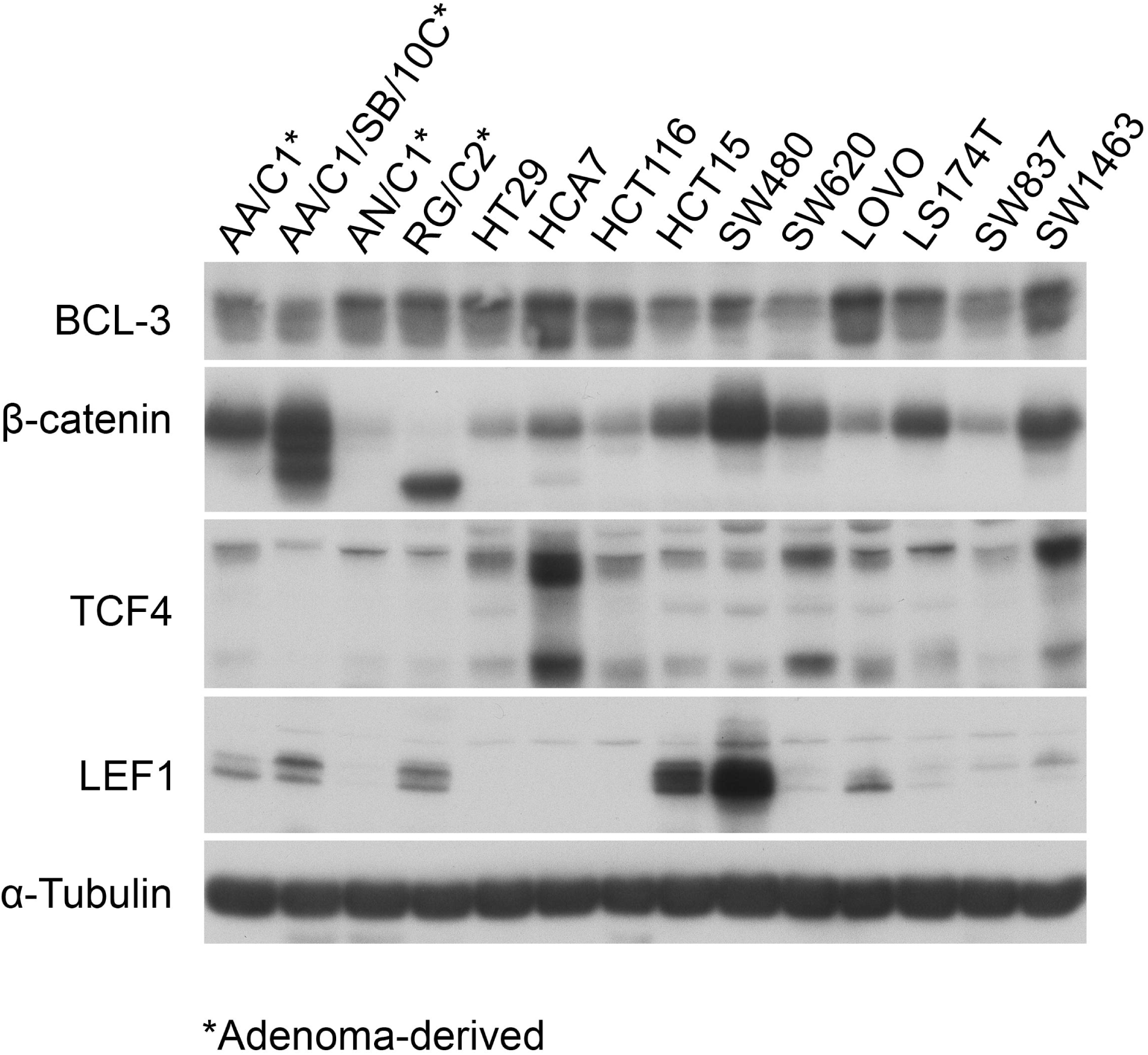
Colorectal adenoma- and carcinoma-derived cell line screen. Western blot analysis of adenoma- and carcinoma-derived colorectal cell lines showing expression of BCL-3 and Wnt transcriptional regulators β-catenin, TCF4 and LEF1. α-Tubulin serves as a loading control.

**Supplementary Figure 2.**
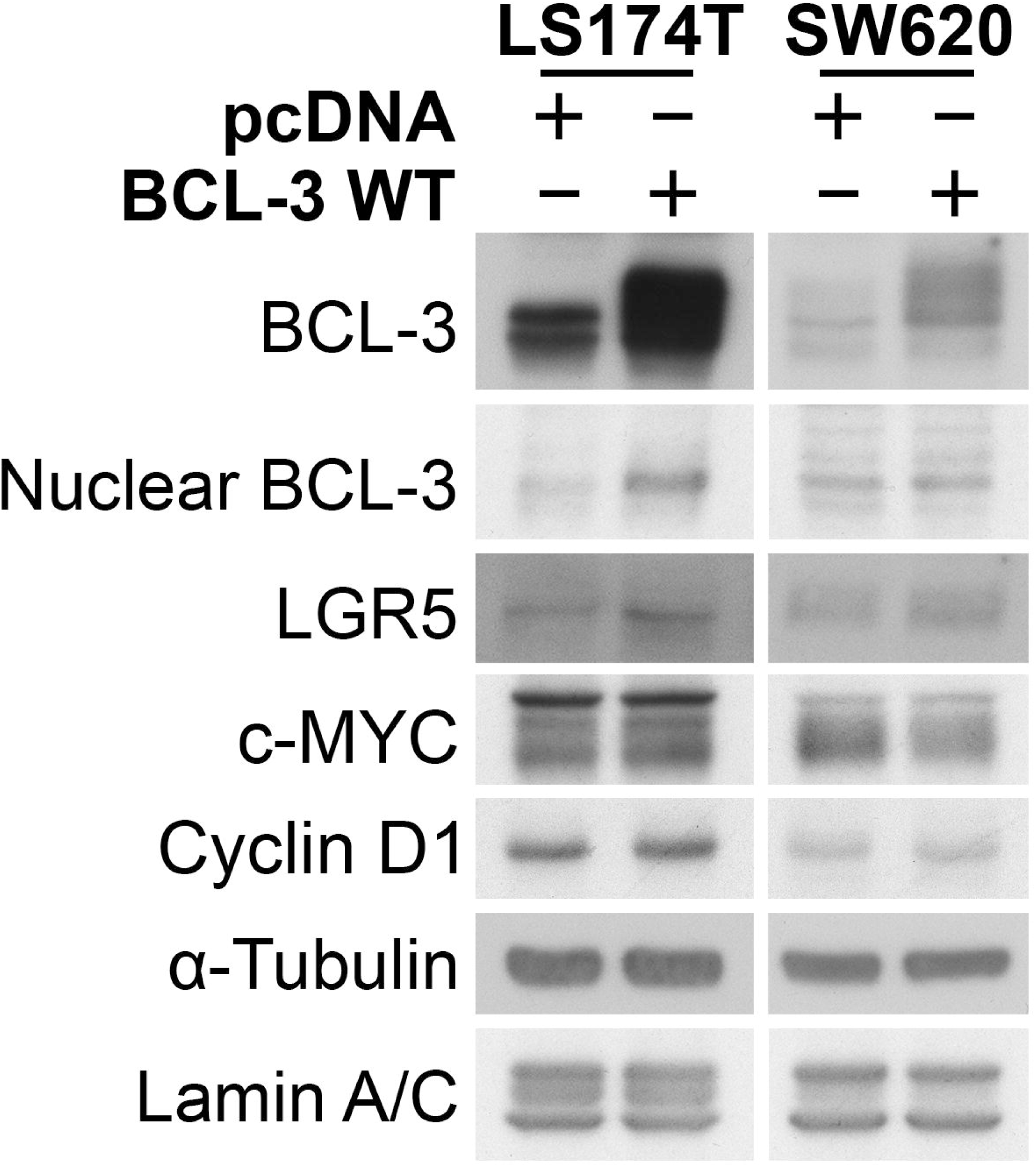
BCL-3 overexpression enhances LGR5 expression. LGR5 expression in LS174T and SW620 cells stably overexpressing BCL-3. LGR5, BCL-3, Cyclin D1 and c-MYC expression was analysed by western blot. α-Tubulin serves as a loading control. Nuclear BCL-3 expression was analysed in nuclear-enriched lysates with Lamin A/C serving as a loading control.

